# SGC-GAK-1: a chemical probe for cyclin G associated kinase (GAK)

**DOI:** 10.1101/376772

**Authors:** Christopher R. M. Asquith, Benedict-Tilman Berger, Jing Wan, James M. Bennett, Michael P. East, Jonathan M. Elkins, Oleg Fedorov, Paulo H. Godoi, Debra M. Hunter, Stefan Knapp, Susanne Müller, Carrow I. Wells, H. Shelton Earp, Timothy M. Willson, William J. Zuercher

**Affiliations:** Structural Genomics Consortium, UNC Eshelman School of Pharmacy, University of North Carolina at Chapel Hill, Chapel Hill, NC 27599, USA; Structural Genomics Consortium, Johann Wolfgang Goethe University, Buchmann Institute for Molecular Life Sciences, Max-von-Laue-Str. 15, D-60438 Frankfurt am Main, DE, Germany; Institute for Pharmaceutical Chemistry, Johann Wolfgang Goethe University, Max-von-Laue-Str. 9, D-60438 Frankfurt am Main, DE, Germany; Department of Medicine, University of North Carolina at Chapel Hill, NC 27599, USA; Lineberger Comprehensive Cancer Center, University of North Carolina at Chapel Hill, NC 27599, USA; Structural Genomics Consortium and Target Discovery Institute, Nuffield Department of Clinical Medicine, University of Oxford, Old Road Campus Research Building, Oxford, 0X3 7DQ, UK; Department of Pharmacology, University of North Carolina at Chapel Hill, NC 27599, USA; Structural Genomics Consortium, Universidade Estadual de Campinas - UNICAMP, Campinas, São Paulo, 13083-886, Brazil

## Abstract

We describe SGC-GAK-1 (**11**), a potent, selective, and cell-active inhibitor of cyclin G associated kinase (GAK), together with a structurally-related negative control SGC-GAK-1N (**14**). SGC-GAK-1 is highly selective in a kinome-wide screen, but cellular engagement assays defined RIPK2 as a collateral target. We identified **18** as a potent inhibitor of RIPK2 lacking GAK activity. Together, the chemical probe set of **11, 14**, and **18** can be used to interrogate the cellular biology of GAK inhibition.

## Introduction

Cyclin G associated kinase (GAK) is a 160 kDa serine/threonine kinase originally identified and named as a direct association partner of cyclin G.^1^ GAK is a member of the numb-associated kinase (NAK) family, which includes AAK1 (adaptor protein 2-associated kinase), STK16/MPSK1 (serine/threonine kinase 16/myristoylated and palmitoylated serine/threonine kinase 1), and BMP2K/BIKE (BMP-2 inducible kinase).^2^ In addition to its kinase domain, the C-terminus of GAK protein has high homology to a domain found in auxilin and tensin.^3^ GAK is ubiquitously expressed and within the cell localizes to the Golgi complex, cytoplasm, and nucleus.^4, 5^

Genetic studies have implicated GAK in several diverse biological processes. Genome-wide association studies have identified single nucleotide polymorphisms in the GAK gene associated with susceptibility to Parkinson’s disease.^6^ GAK like other members of the NAK family is involved in membrane trafficking and sorting of proteins and as an essential cofactor for HSC70-dependent uncoating of clathrin-coated vesicles in the cytoplasm.^7, 8^ Through its association with cyclin G, GAK is also required for maintenance of centrosome maturation and progression through mitosis.^9^ GAK is over-expressed in osteosarcoma cell lines and tissues where it contributes to proliferation and survival.^10^ Notably, GAK expression increases during prostate cancer progression to androgen independence and is positively correlated with the Gleason score in surgical specimens from prostate cancer patients.^11, 12^

We have embarked on a program to develop small molecule chemical probes to elucidate the biological role of kinases such as GAK that belong the lesser-studied portion of the kinome.^13^ To be useful research reagents, these chemical probes must be potent, selective, and cell-active.^14^ Specifically the aspirational criteria we are following are (i) *in vitro* biochemical IC_50_ < 50 nM; (ii) more than 30-fold selective relative to other kinases in a large assay panel such as DiscoverX KINOMEscan^®^; and (iii) cellular activity or target engagement IC_50_ < 1 μM. In addition, the probe should be accompanied by a negative control compound that is structurally related but at least several orders of magnitude less potent at the primary kinase target.^14^

A number of quinazoline-based clinical kinase inhibitors show cross reactivity with GAK, including the approved drugs gefitinib, erlotinib, and bosutinib (**Figure 1**).^15^ These drugs were designed as inhibitors of either epidermal growth factor receptor (EGFR) or SRC kinase and show similar or higher affinity for a variety other kinases, making them ineffective tools for studying the biology of GAK.^16^ It is not known whether the clinical efficacy or adverse events observed with these kinase inhibitor drugs are associated with their cross-activity on GAK. Notably, it has been proposed that GAK inhibition causes clinical toxicity due to pulmonary alveolar dysfunction. This hypothesis is based in part on the phenotype of transgenic mice expressing catalytically inactive GAK but has not yet been tested with a selective small molecule GAK inhibitor.^17^

**Figure 1.**
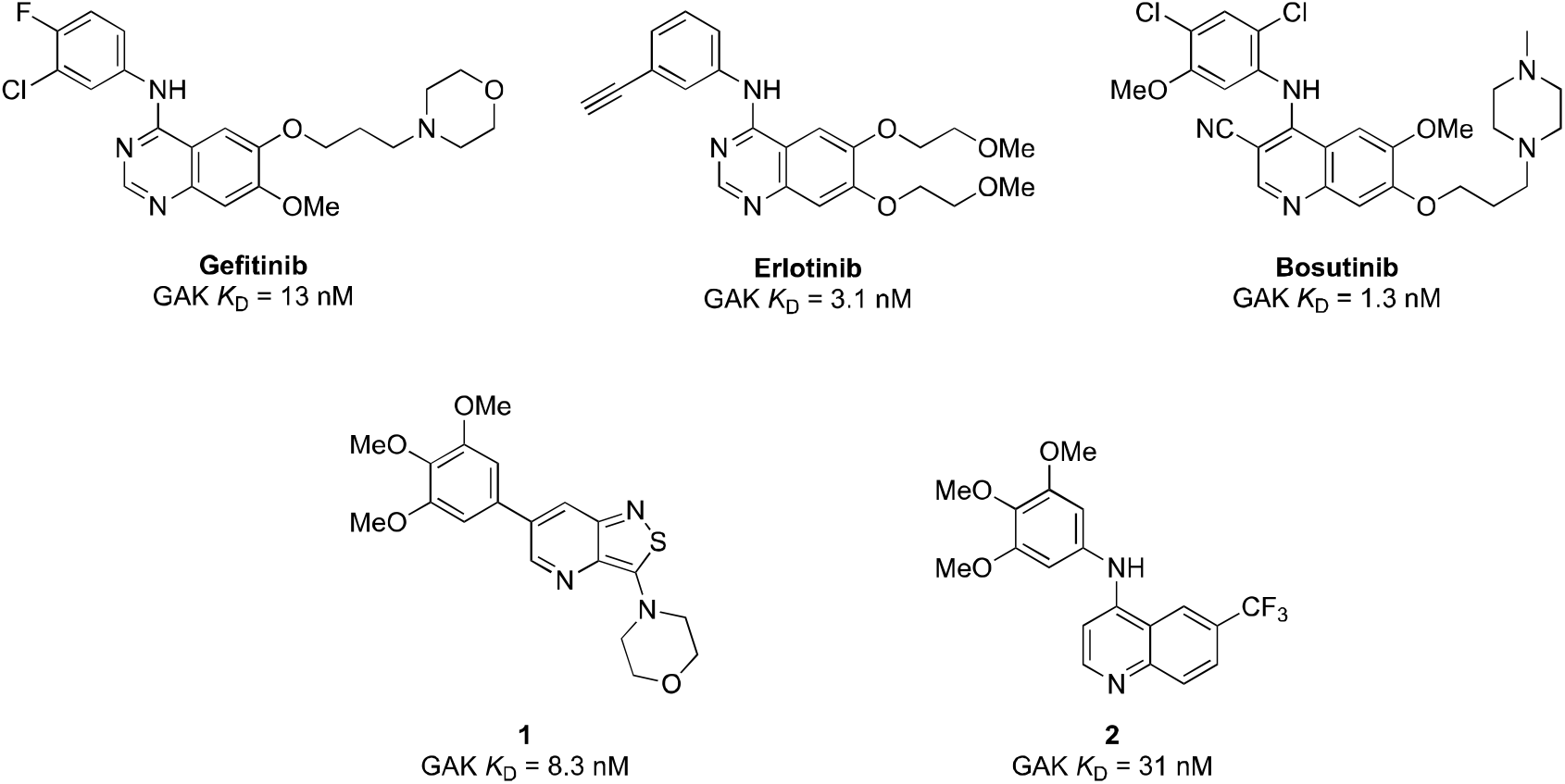
Previously described inhibitors of GAK.

Isothiazolo[5,4-*b*]pyridine **1** has been described as a selective inhibitor of GAK.^18^ However, **1** shows cross reactivity with other kinases, including KIT, PDGFRB, FLT3, and MEK5, any of which could lead to confounding biology in cells. The availability of a chemical probe with improved selectivity for GAK, increased cellular potency, and a narrower spectrum of off-targets would be a valuable tool to probe GAK biology. We previously described the identification of a series of 4-anilinoquinoline inhibitors of GAK, exemplified by **2**.^19^ The optimization of this series to a potent, selective, and cell-active chemical probe is described.

## Results

Quinoline **2** was initially profiled at 1 μM across an assay panel of over 400 wild-type human kinases. Subsequent *K*_D_ determination for those kinases with > 60% binding at 1 μM identified three kinases (receptor-interacting protein kinase 2, RIPK2; AarF domain containing kinase 3, ADCK3; and nemo-like kinase, NLK) with *K*_D_ values within 30-fold of that of GAK (**Table S4**). While little is known about the activity of quinazolines on ADCK3, and NLK, a prior study of quinazoline inhibitors of RIPK2 showed a hydrogen bonding interaction between Ser25 of RIPK2 near the solvent exposed portion of the ATP-binding pocket.^20^ Since GAK has Ala47 as the corresponding residue, we hypothesized that small, nonpolar substituents at the quinoline 6- and 7-positions held the potential for improving the selectivity over RIPK2.

To investigate this hypothesis and define the SAR against ADCK and NLK, we profiled several quinolines prepared in our previous study against the three collateral targets in heterogeneous assays at DiscoverX involving displacement of kinase protein from an immobilized inhibitor (**Table 1**).^19^ 6,7-Dimethoxyquinoline **3** had the highest GAK affinity but also showed increased binding to RIPK2 relative to **2**. Replacement of either methoxy group with a hydrogen, as in **4** and **5**, led to reduced affinity on all four kinases. Notably, however, the unsubstituted quinoline **6** showed an improved selectivity profile, although it had the poorest affinity for GAK among the analogs. The nitrile isomers **7** and **8** both retained single digit nanomolar affinity for GAK yet had different effects on selectivity profile: the 6-isomer (**7**) had an improved selectivity profile while the 7-isomer (**8**) showed binding to all three off-targets within ten-fold. Addition of a tertiary butyl group in the 6-position (**9**) led to improvement of the selectivity profile against RIPK2 and ADCK3 with a decrease in selectivity over NLK compared with **2**.

**Table 1.**
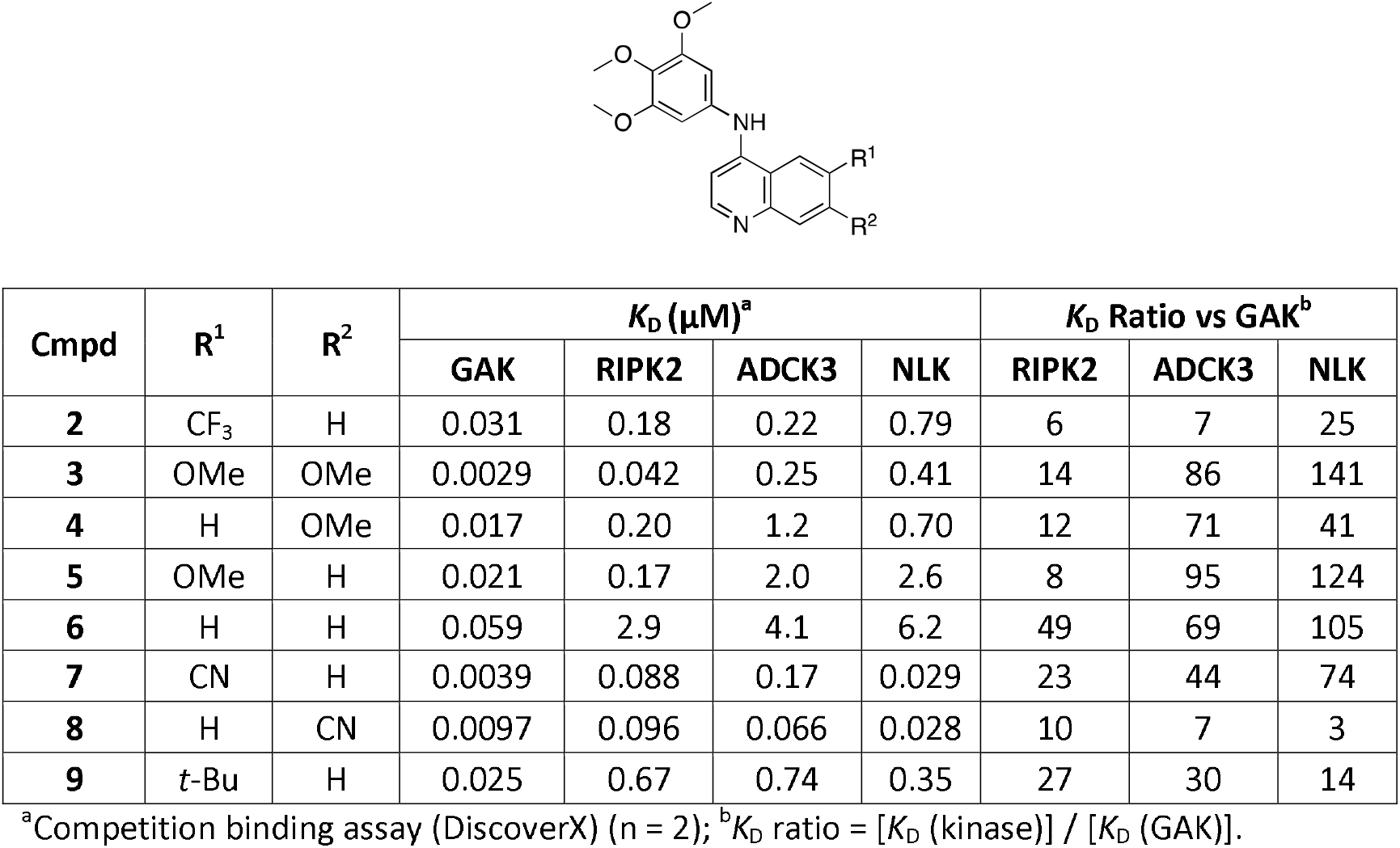
Affinity of 4-anilinoquinolines for GAK, RIPK2, ADCK3 and NLK.

| Cmpd | R<sup>1</sup> | R<sup>2</sup> | K<sub>D</sub> (μM)<sup>a</sup> |  |  |  | K<sub>D</sub> Ratio vs GAK<sup>b</sup> |  |  |
| --- | --- | --- | --- | --- | --- | --- | --- | --- | --- |
|  |  |  | GAK | RIPK2 | ADCK3 | NLK | RIPK2 | ADCK3 | NLK |
| **2** | CF<sub>3</sub> | H | 0.031 | 0.18 | 0.22 | 0.79 | 6 | 7 | 25 |
| **3** | OMe | OMe | 0.0029 | 0.042 | 0.25 | 0.41 | 14 | 86 | 141 |
| **4** | H | OMe | 0.017 | 0.20 | 1.2 | 0.70 | 12 | 71 | 41 |
| **5** | OMe | H | 0.021 | 0.17 | 2.0 | 2.6 | 8 | 95 | 124 |
| **6** | H | H | 0.059 | 2.9 | 4.1 | 6.2 | 49 | 69 | 105 |
| **7** | CN | H | 0.0039 | 0.088 | 0.17 | 0.029 | 23 | 44 | 74 |
| **8** | H | CN | 0.0097 | 0.096 | 0.066 | 0.028 | 10 | 7 | 3 |
| **9** | *t*-Bu | H | 0.025 | 0.67 | 0.74 | 0.35 | 27 | 30 | 14 |

<sup>a</sup>Competition binding assay (DiscoverX) (n = 2); <sup>b</sup>K<sub>D</sub> ratio = [K<sub>D</sub> (kinase)] / [K<sub>D</sub> (GAK)].

Comparison of the two matched molecular pairs with methoxy and nitrile (**4** and **8**; **5** and **7** respectively) suggested that electron withdrawing substituents at the quinoline 6-position should be investigated further. The 6-halogenated quinolines **10-12** were prepared as described previously by nucleophilic substitution of the corresponding 4-chloroquinolines with 3,4,5-trimethoxyaniline.^19^ When profiled in the panel of the four kinases, these halogen-substituted analogs (**10-12**) showed high GAK affinity, all with *K*_D_ < 10 nM (**Table 2**). Analog **11** also displayed remarkable selectivity, with the closest off-target kinase (RIPK2) being > 50-fold weaker in affinity.

**Table 2.** Screening results of GAK, RIPK2, ADCK3 and NLK on the 4-anilinoquinoline scaffold.

| Cmpd | R | $K_D$ (μM)<sup>a</sup> | | | | $K_D$ Ratio<sup>b</sup> vs GAK | | |
| --- | --- | --- | --- | --- | --- | --- | --- | --- |
|  |  | GAK | RIPK2 | ADCK3 | NLK | RIPK2 | ADCK3 | NLK |
| **10** | Cl | 0.0067 | 0.085 | 0.17 | 0.65 | 13 | 25 | 97 |
| **11** | Br | 0.0019 | 0.11 | 0.19 | 0.52 | 58 | 100 | 274 |
| **12** | I | 0.0079 | 0.069 | 0.19 | 0.66 | 9 | 24 | 84 |

<sup>a</sup>Competition binding assay (DiscoverX) (n = 2) ; <sup>b</sup> $K_D$ ratio = [ $K_D$ (kinase)] / [ $K_D$ (GAK)].

To confirm the selectivity profiles within the NAK subfamily, the affinity of **11** was evaluated for for GAK, AAK1, BMP2K, and STK16 in time-resolved fluorescence energy transfer (TR-FRET) ligand binding displacement assays (**Figure 2A**). This homogeneous assay system is distinct from the heterogeneous configuration of the kinase binding assays at DiscoverX. **11** displayed an impressive profile of over 16000-fold selectivity relative to each of the other three NAK family members. In addition to the NAK family binding assay panel, a differential scanning fluorimetry (DSF) assay^21^ and isothermal calorimetry (ITC) were employed to confirm direct interaction with GAK (**Figure 2B** and **2C**). **11** induced a significant increase in thermal stability of the isolated GAK kinase domain, suggesting strong binding of the inhibitor. These data were confirmed by ITC which revealed potent affinity for GAK (*K*_D_ = 4.5 nM).

**Figure 2.**
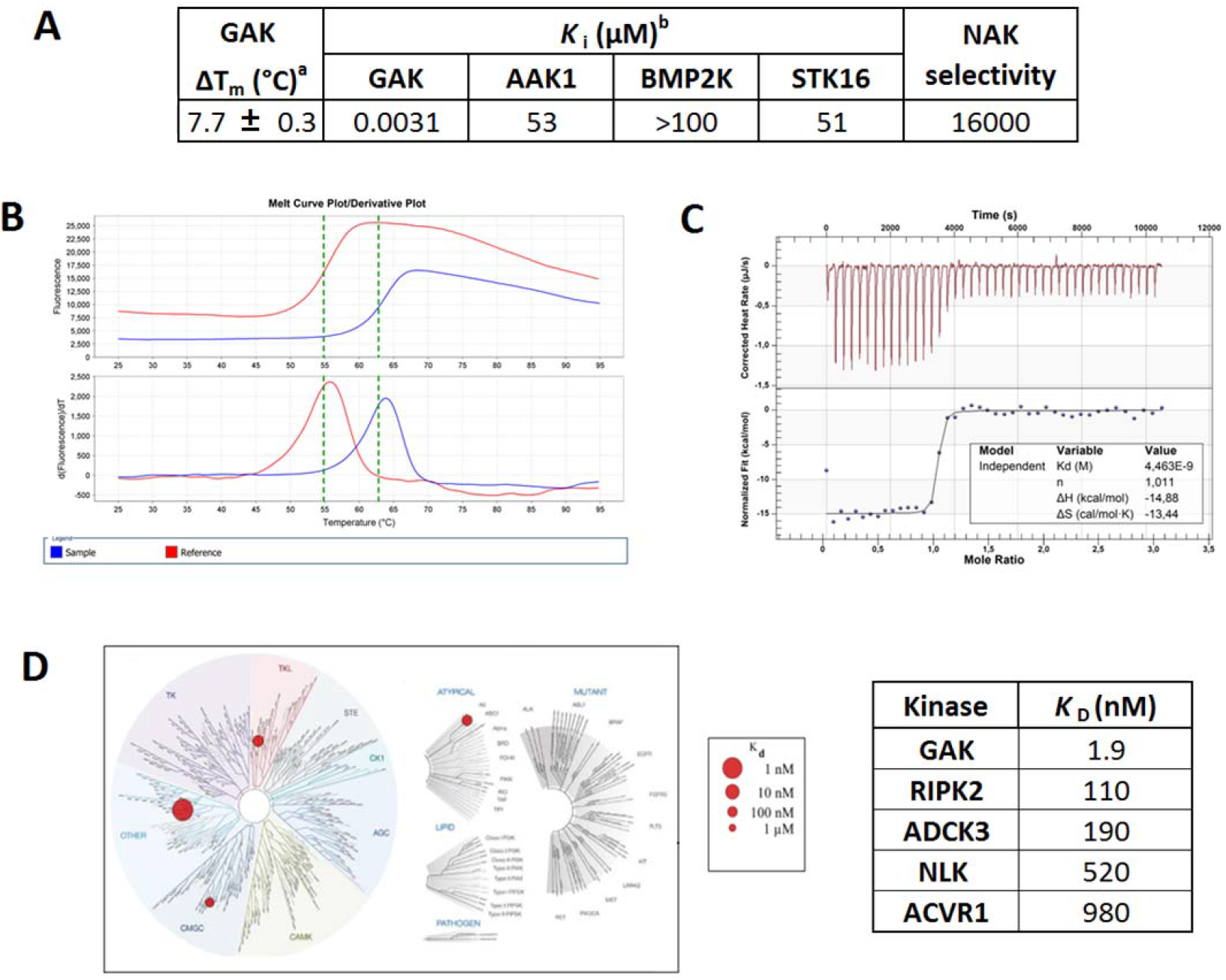
(A) Binding affinity of **11** for NAK family kinases.^a^Shift in temperature of GAK thermal denaturation as measured by differential scanning fluorimetry (n = 4). ^b^TR-FRET-based ligand binding displacement assay (n = 1); (B) DSF curve of GAK and compound **11;** (C) ITC of compound **11** on GAK; (D) KINOMEscan^®^ results and select *K*_D_ determinations for compound **11**.

Given the encouraging selectivity profile of **11**, we assessed its selectivity in the KINOMEscan^®^ assay panel (**Figure 2D**). *K*_D_ determinations were obtained on all kinases with binding greater than 50% at 1 μM (**Table S2**). **11** showed >50-fold selectivity relative to all 400 kinases. The closest off-targets were RIPK2, ADCK3, NLK and ACVR1 (**Figure 2** and **Table S5**).

An established best practice for the use of chemical probes in biological assays is to test in parallel a compound closely related in structure to the probe but inactive for the target protein.^14^ Demonstration of divergent results between the chemical probe and its negative control increases confidence that the engagement of the target protein is responsible for the observed biology. We designed and prepared two potential negative control compounds that were anticipated to lack GAK activity. First, **13** incorporated a 2-methylquinoline that would directly hinder hydrogen bonding to the hinge region of the kinase. Second, the *N*-methyl compound **14** was expected to disrupt a water network necessary for favorable compound binding (Asquith, C.R.M. and Laitinen, T., unpublished results). Neither compound induced thermal stabilization of GAK protein, and both demonstrated >10,000-fold weaker GAK binding affinity in the TR-FRET assay (**Table 4**). The KINOMEscan^®^ and subsequent *K*_D_ determinations of **13** and **14** identified only a single kinase with submicromolar affinity (**14**: interleukin 1 receptor associated kinase 3, IRAK3 *K*_D_ = 740 nM) (**Table S3**). Thus, both compounds were suitable as GAK negative controls. We selected **14** as the negative control based on its higher selectivity in the NAK family ligand binding displacement assay. An additional advantage of **14** is that the subtle structural modification contributing to loss of GAK activity was not expected to directly perturb the important kinase hinge binding interaction.

**Table 4.** NAK family binding assay results of potential negative control compounds **13** and **14**.

| Cmpd | R<sup>1</sup> | R<sup>2</sup> | GAK<br>$\Delta T_m$ (°C)<sup>a</sup> | $K_i$ (μM)<sup>b</sup> | | | |
| --- | --- | --- | --- | --- | --- | --- | --- |
|  |  |  |  | AAK1 | BMP2K | GAK | STK16 |
| **13** | Me | H | -0.52 | >100 | >100 | 19 | >100 |
| **14** | H | Me | -0.30 | 78 | 95 | 2.5 | 15 |

<sup>a</sup>Shift in temperature of thermal denaturation of GAK as measured by differential scanning fluorimetry; <sup>b</sup>TR-FRET-based ligand binding displacement assay (n = 1).

Having confirmed potent GAK binding and selectivity relative to other NAK family members, we next sought to demonstrate that activity and selectivity of **11** in live cells using a NanoBRET target engagement assay.^22^ Compounds **11** and **14** were evaluated for their ability to displace a fluorescent tracer molecule from the ATP binding site of a transiently expressed full length GAK protein *N*-terminally fused with NanoLuc luciferase (Nluc) in HEK293T cells. In the absence of compound, the tracer molecule and fusion protein were in proximity and able to generate an observable bioluminescence resonance energy transfer (BRET) signal. Increasing amounts of added compound led to displacement of the tracer and concomitant loss of BRET signal. 6-Bromoquinoline **11** had high affinity for GAK in cells (IC_50_ = 120 nM), although the measured IC_50_ was 24-fold higher than the *in vitro K_D_* value. We interpret the difference as most likely due to the effect of competition with ATP in cells. The negative control compound **14** showed no cellular affinity for GAK up to a concentration of 5.0 μM. Notably, the previously reported GAK inhibitor 1 displayed GAK cellular potency in the single digit μM range, approximately ten-fold weaker than **11**. We also evaluated **11** and **14** in a RIPK2 NanoBRET assay as this kinase was the closest off-target. We were surprised to find that **11** had only three-fold selectivity for GAK over RIPK2 in cells (**Figure 3**, GAK IC_50_ =120 ± 50 nM; RIPK2 IC_50_ = 360 ± 190 nM). Negative control **14** did not bind RIPK2 in cells at concentrations up to 10 μM. The previously described GAK inhibitor 1 also showed no affinity for RIPK2.

**Figure 3.**
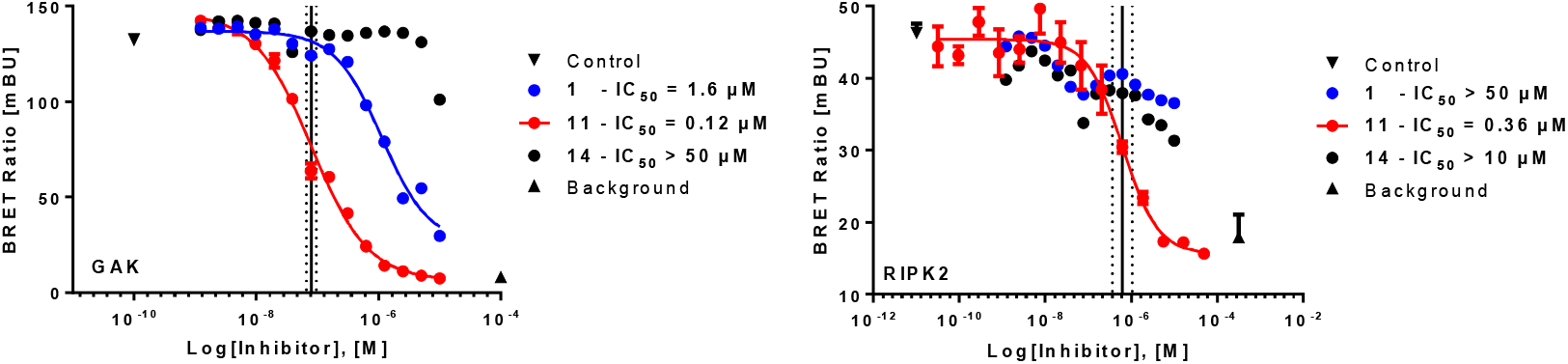
Evaluation of **1, 11** and **14** in GAK and RIPK2 NanoBRET cellular target engagement assays. A measurement where the inhibitor was substituted with DMSO is referred to as ‘control’ and a measurement where both tracer and inhibitor were substituted by DMSO is referred to as ‘background’. The corresponding IC_50_ is shown as the average of two individual experiments of duplicates from which the standard error of the mean (SEM) was calculated.

Given the observed cross-activity of **11** with RIPK2 in cells, we sought to identify an additional control compound with activity on RIPK2 but not GAK. Surveying known RIPK2 inhibitors (**Table 5**), we initially selected OD36 (**15**), since it had previously been characterized as selective for RIPK2 across a panel of over 400 wild type human kinase assays (although this panel did not include GAK). We also expected that the distinct macrocyclic chemotype of **15** would have a lower likelihood of showing affinity for GAK relative to a quinoline.^23^ However, we were surprised to observe that **15** was nearly equipotent at engagement of both RIPK2 and GAK in cells. Affinity of **15** for GAK was confirmed in the TR-FRET *in vitro* binding assay. Next, we selected three quinoline-based RIPK2 inhibitors (**16-18**) previously reported by scientists at GlaxoSmithKline.^20, 24^ Disappointingly, indazole-quinazoline **16** and the modified pyrazole-quinazoline **17** showed high affinity for GAK, in the TR-FRET assay. Fortunately, benzothiazole-quinazoline **18**, showed low affinity for GAK in the TR-FRET assay and in the NanoBRET assay.^25^ **18** demonstrated potent engagement with RIPK2 in cells and was selected to control for RIPK2 activity.

**Table 5.**
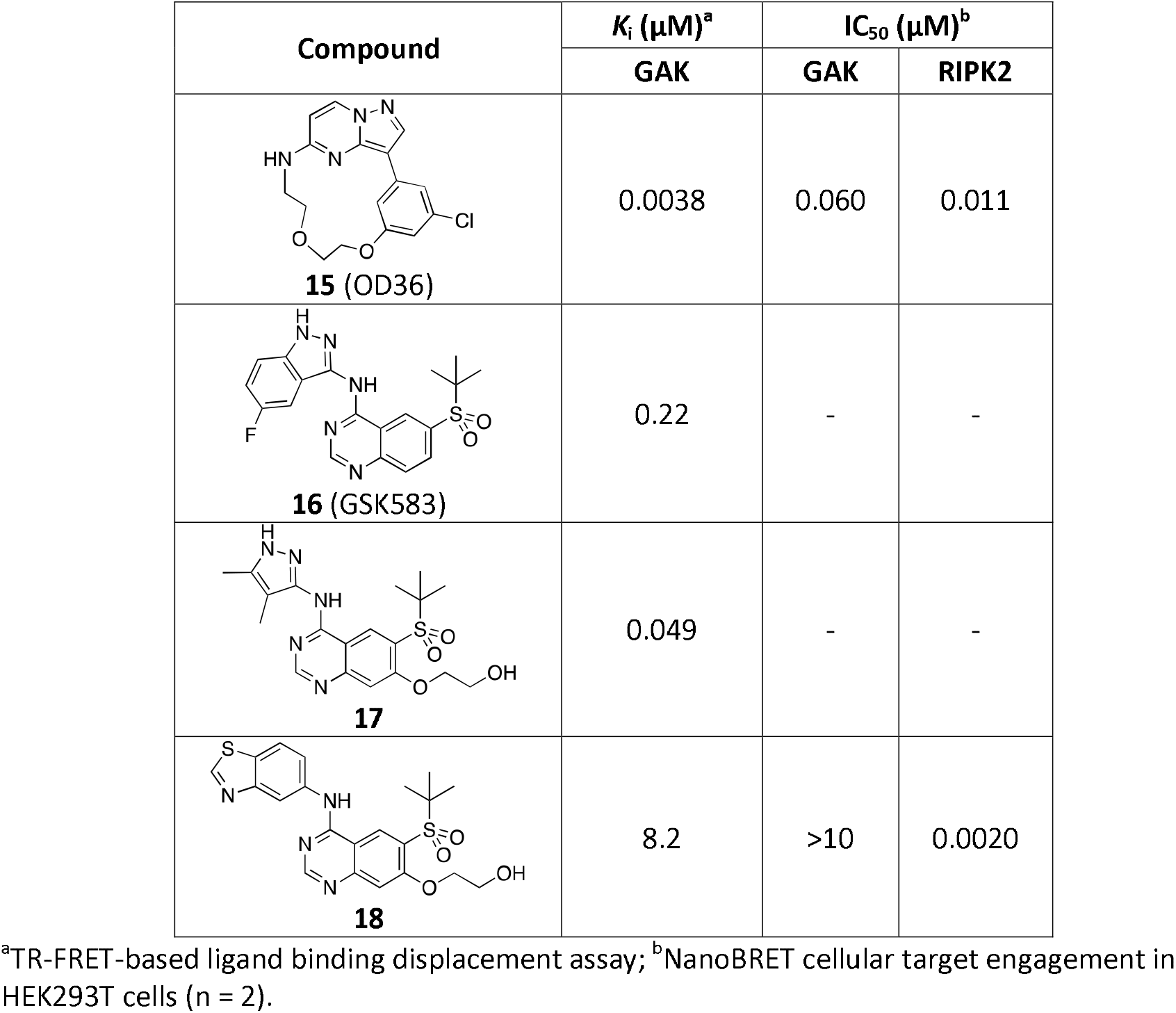
Evaluation of potential RIPK2 control compounds.

We propose that, when used in parallel, SGC-GAK-1 (**11**) and the controls **14** and **18** can be used as a probe set to study the biology of GAK in cells. For example, to address the proposed role of GAK in prostate cancer,^12^ the probe set was tested for the effect on the viability of five well-characterized prostate cancer cell lines (PC3, DU145, LNCaP, VCaP, and 22Rv1 cells). Western blotting experiments confirmed that GAK was expressed in these cell lines (**Figure S5**). In an initial experiment, the cell lines were incubated with **11, 14**, and **18** at three different concentrations (0.1, 1.0, and 10 μM), and viability was measured after either 48 h or 72 h. No effect was observed with the control compounds **14** and **18** in any of the cell lines at these concentrations. However, SGC-GAK-1 (**11**) showed growth inhibition in all cell lines at 10 μM, albeit minimally in PC3 (12 %) and DU145 (21 %) cells but much stronger in LNCaP (53 %), VCaP (64 %), and 22Rv1 (61 %) cells. IC_50_ values for **11** were then determined for these latter three cell lines in biological triplicate measurements (**Table 6** and **Figure 4**). **11** showed modest antiproliferative activity in LNCaP cells but greatly increased potency in VCaP and 22Rv1 cells. Overall, these results demonstrate selective effects of the GAK chemical probe in a subset of prostate cancer cell lines.

**Figure 4.**
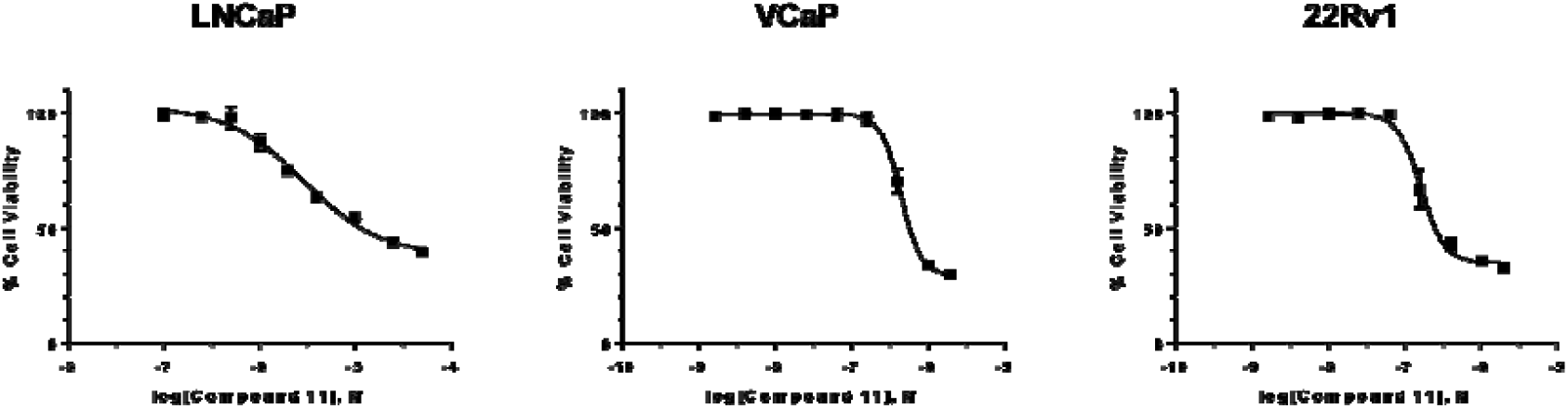
Representative dose response measurements for **11** in LNCaP, 22Rv1, and VCaP cell lines.

**Table 6.**
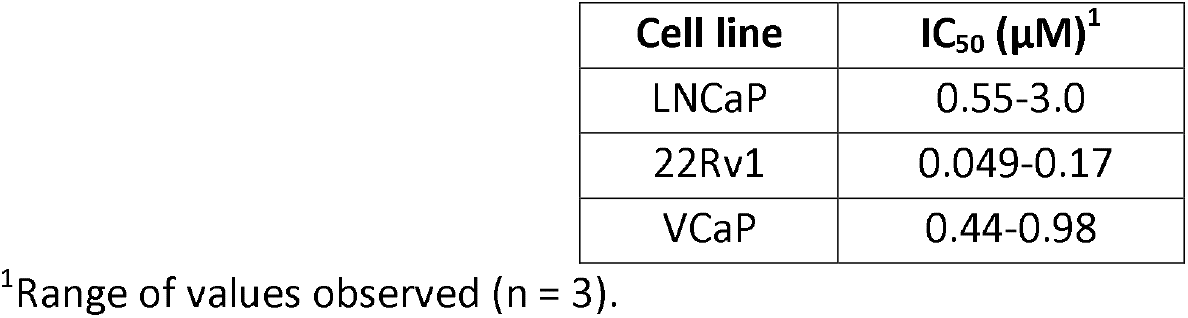
Effects of **11** on the viability of prostate cancer cell lines at 48 h (LNCaP and 22Rv1) or for 72 h (VCaP) after compound treatment.

Previous reports have demonstrated that GAK is required for proper mitotic progression. Specifically, siRNA knockdown of GAK in HeLa cells was reported to induce cell cycle arrest at the metaphase and increase levels of phosphorylated histone H3 Ser10.^26^ Prolonged metaphase arrest is known to lead to activation of apoptotic pathways.^27^ We observed that treatment of 22Rv1 cells with **11** led to PARP cleavage, a marker of cells undergoing apoptosis (**Figure S6**), and an increase in phosphorylated histone H3 Ser10, as had been observed in with GAK targeting siRNA.^26^ The present results demonstrate similar effects through pharmacological inhibition of the GAK kinase domain as has been reported for siRNA mediated depletion the GAK protein.

## Discussion

4-Anilinoquinolines and quinazolines are common structural templates for kinase inhibitors, including several approved medicines, with many more under clinical investigation.^16^ We previously identified a series of 4-anilinoquinolines as narrow spectrum inhibitors of GAK and now describe our efforts leading to the potent, selective, and cell-penetrant GAK probe compound **11**. GAK chemical probe **11** offers several advantages over the previously reported isothiazolo[4,3-*b*]pyridine GAK inhibitor 1, including superior cellular potency, improved kinase selectivity. Furthermore, **11** is accompanied by control compounds **14** and **18** that when tested in parallel result permit assignment of the on-target effects of the GAK chemical probe. This GAK probe and control compounds are openly available, and we expect they will find utility in the elucidation of GAK biology in cells.

The identification of RIPK2 as a cellular cross-target of **11** highlights the importance of compound profiling in multiple assay systems. Chemical probe **11** demonstrated a > 50-fold difference in affinity between GAK and RIPK2 in heterogeneous, ATP-free binding assays. In contrast, the compound was nearly equipotent (within 4-fold affinity) on these two kinases when evaluated in the NanoBRET live cell target engagement assay. The NanoBRET assay has been shown to closely recapitulate conditions encountered in cell-based phenotypic screens, using full length protein and being measured at physiological levels of ATP and the presence of endogenous protein partners.^22^ To support robust interpretation of the biology of **11**, we identified the quinazoline **18** as a RIPK2 inhibitor that lacks affinity for GAK as means to control for this off-target effect in cells.

The observation that GAK expression is associated with prostate cancer progression prompted us to explore the effect of GAK inhibition across a range of cell lines. To date all studies of the role of GAK in prostate cancer have utilized knock-down of the whole protein.^28^ We now show for the first time that inhibition of the GAK kinase domain alone can block the growth of prostate cancer cells. Our results demonstrate that the kinase and not the auxilin-homology domain is critical for this phenotype in cells. SGC-GAK-1 (**11**), but neither of the control compounds **14** nor **18**, blocked growth of 22Rv1, LNCaP, and VCaP cell lines but not DU145 or PC3 cell lines. Since neither DU145 nor PC3 express the AR, their proliferation is not expected to be AR-dependent. In contrast LNCaP, 22Rv1 and VCaP are all AR positive and are cell models for androgen-dependent growth. Interestingly, the most potently inhibited cell lines (VCaP and 22Rv1) express AR splice variants including AR-V7. These splice variants are often increased in patients who become resistant to androgen deprivation.^29^ The response to **11** in the LNCaP cell line which expresses wild-type AR but not AR-V7 was less than that of the two splice variant expressing lines. Why the AR-V7 expressing lines would be more sensitive and what are the mechanisms by which GAK inhibitors target a subset of AR expressing prostate cancer cells are the subjects of on-going investigation.

In summary, we have generated the first chemical probe set for the poorly studied kinase GAK. SGC-GAK-1, when used in parallel with **14** and **18**, allows assignment of the biology of GAK kinase inhibition in cells. Using this probe set we demonstrate that the GAK kinase domain is a potential target for inhibition of prostate cancer growth in cells that express AR splice variants associated with poor clinical prognosis.

## Acknowledgment

The SGC is a registered charity (number 1097737) that receives funds from AbbVie, Bayer Pharma AG, Boehringer Ingelheim, Canada Foundation for Innovation, Eshelman Institute for Innovation, Genome Canada, Innovative Medicines Initiative (EU/EFPIA) [ULTRA-DD grant no. 115766], Janssen, Merck KGaA Darmstadt Germany, MSD, Novartis Pharma AG, Ontario Ministry of Economic Development and Innovation, Pfizer, São Paulo Research Foundation-FAPESP, Takeda, and Wellcome [106169/ZZ14/Z]. Antti Poso and Tuomo Laitinen (University of Eastern Finland), David H. Drewry (SGC-UNC), and Stephen J. Capuzzi (UNC-CH) are thanked for informative discussions. We also thank Dr. Brandie Ehrmann for LC-MS/HRMS support provided by the Mass Spectrometry Core Laboratory at the University of North Carolina at Chapel Hill.

## Experimental

All reactions were performed using flame-dried round-bottomed flasks or reaction vessels unless otherwise stated. Where appropriate, reactions were carried out under a nitrogen atmosphere with dry solvents, unless otherwise stated. Yields refer to chromatographically and spectroscopically pure isolated yields. Reagents were purchased at the highest commercial quality and used without further purification, unless otherwise stated. Reactions were monitored by thin-layer chromatography carried out on 0.25 mm E. Merck silica gel plates (60_F-254_) using ultraviolet light as visualizing agent. NMR spectra were recorded on a Varian Inova 400 or Inova 500 spectrometer and were calibrated using residual protic solvent as an internal reference (CDCI_3_:^1^H NMR = 7.26, ^13^C NMR = 77.16). The following abbreviations or combinations thereof were used to explain the multiplicities observed: s = singlet, d = doublet, t = triplet, q = quartet, m = multiplet, br = broad. Liquid chromatography (LC) and high-resolution mass spectra (HRMS) were recorded on a ThermoFisher hybrid LTQ FT (ICR 7T). Compounds **2-14** were synthesised as previously reported, and compounds **2-9** were consistent with previous reports.^19^

### General procedure for the synthesis of 4-anilinoquinolines

6-Bromo-4-chloroquinoline (1.0 eq.), 3,4,5-trimethoxyaniline (1.1 eq.) and DIPEA (2.5 eq.) were suspended in ethanol (30 mL) and refluxed at 90 °C for 18 h. The crude mixture was purified by flash chromatography ethyl acetate:hexane followed by 1-5 % methanol:ethyl acetate and solvent removed under reduced pressure to yield the product as a free following solid.

6-Chloro-*N*-(3,4,5-trimethoxyphenyl)quinolin-4-amine (**10**) as a mustard solid (198 mg, 0.58 mmol, 38%) MP 161-163 °C; ^1^H NMR (400 MHz, DMSO-*d*_6_) δ 10.90 (s, 1H), 8.99 (d, *J* = 2.1 Hz, 1H), 8.49 (d, *J* = 6.9 Hz, 1H), 8.14 (d, *J* = 9.1 Hz, 1H), 8.03 (dd, *J* = 9.0, 2.2 Hz, 1H), 6.93 (d, *J* = 6.9 Hz, 1H), 6.81 (s, 2H), 3.80 (s, 6H), 3.72 (s, 3H). ^13^C NMR (101 MHz, DMSO-*d*_6_) δ 153.8, 153.6 (2C), 143.5, 137.8, 136.4, 133.6, 132.9, 131.3, 123.1, 122.9, 118.2, 102.9 (2C), 100.9, 60.2, 56.1 (2C). HRMS-ESI (m/z): [M+H]^+^ calcd for C_18_H_18_N_2_O_3_Cl: 345.1006, found 345.0993; LC - T_r_ = 3.56 min, purity >98 %.

6-Bromo-*N*-(3,4,5-trimethoxyphenyl)quinolin-4-amine (**11**) as a mustard solid (308 mg, 0.79 mmol, 64%) MP 178-180 °C;^1^H NMR (400 MHz, DMSO-*d*_6_) δ 11.06 (s, 1H), 9.19 (d, *J* = 1.7 Hz, 1H), 8.48 (d, *J* = 6.9 Hz, 1H), 8.32 – 7.82 (m, 2H), 6.92 (d, *J* = 6.9 Hz, 1H), 6.82 (s, 2H), 3.80 (s, 6H), 3.72 (s, 3H); ^13^C NMR (101 MHz, DMSO-*d*_6_) δ 154.0, 153.6 (2C), 143.0, 137.6, 136.5, 136.3, 132.8, 126.2, 122.7, 119.7, 118.5, 103.0 (2C), 100.9, 60.2, 56.1 (2C). LC Chromatogram HRMS-ESI (m/z): [M+H]^+^ calcd for C_18_H_18_N_2_O_3_Br: 389.0500, found 389.0490; LC - T_r_ = 3.60 min, purity >98 %.

6-lodo-*N*-(3,4,5-trimethoxyphenyl)quinolin-4-amine (**12**) as a bright yellow solid (321 mg, 0.74 mmol, 71%) MP 172-175 °C; ^1^H NMR (400 MHz, DMSO-*d*_6_) δ 10.61 (s, 1H), 9.19 (s, 1H), 8.46 (d, *J* = 6.6 Hz, 1H), 8.19 (d, *J* = 10.2 Hz, 1H), 7.87 (d, *J* = 8.8 Hz, 1H), 6.92 (d, *J* = 6.5 Hz, 1H), 6.78 (s, 2H), 3.79 (s, 6H), 3.71 (s, 3H); ^13^C NMR (101 MHz, DMSO-*d*_6_) δ 153.5 (2C), 152.3, 144.6, 140.8, 139.8, 136.0, 133.5, 131.8, 124.0, 119.4, 102.5 (2C), 101.1, 92.2, 60.2, 56.1 (2C). LC Chromatogram HRMS-ESI (m/z): [M+H]^+^ calcd for C_18_H_18_N_2_O_3_I: 437.0362; found 437.0344; LC - T_r_ = 3.73 min, purity >98 %.

*N*-Methyl-6-(trifluoromethyl)-N-(3,4,5-trimethoxyphenyl)quinolin-4-amine (**13**) (half reaction size) as a colourless solid (114 mg, 0.29 mmol, 45 %) MP 106-108 °C; ^1^H NMR (400 MHz, DMSO-*d*_6_) δ 8.85 (d, *J* = 5.2 Hz, 1H), 8.11 – 8.05 (m, 1H), 7.85 – 7.77 (m, 2H), 7.25 (d, *J* = 5.2 Hz, 1H), 6.39 (s, 2H), 3.61 (s, 3H), 3.59 (s, 6H), 3.48 (s, 3H); ^13^C NMR (101 MHz, DMSO-*d*_6_) δ 153.6, 153.4, 153.0, 150.5, 145.6, 134.8, 131.1, 125.4, 124.4, 124.1, 123.9 (q, *J* = 2.9 Hz), 123.1 (q, *J* = 4.6 Hz), 122.7, 121.3, 111.3, 101.3, 60.1, 56.1 (2C), 42.8. HRMS-ESI (m/z): [M+H]^+^ calcd for C_20_H_20_N_2_O_3_F_3_: 393.1426, found 393.1413; LC - T_r_ = 4.17 min, purity >98 %.

2-Methyl-6-(trifluoromethyl)-N-(3,4,5-trimethoxyphenyl)quinolin-4-amine (**14**) as a yellow solid (330 mg, 0.84 mmol, 69%) MP 240 °C decomp.; NMR (400 MHz, DMSO-*d*_6_) δ 10.87 (s, 1H), 9.20 (s, 1H), 8.23 (s, 2H), 6.87 (s, 1H), 6.79 (s, 2H), 3.81 (s, 6H), 3.73 (s, 3H), 2.65 (s, 3H); ^13^C NMR (101 MHz, DMSO-*d*_6_) δ 157.3, 154.7, 154.1 (2C), 136.9, 133.2, 129.2-129.1 (m, 1C), 126.7, 126.7, 126.4, 125.7, 123.0, 122.4 (d, *J* = 3.8 Hz), 116.2, 103.4, 102.0, 60.6, 56.6 (2C), 20.9. LC Chromatogram; HRMS-ESI (m/z): [M+H]^+^ calcd for C_20_H_19_F_3_N_2_O_3_: 393.1426, found 393.1413; LC - T_r_ = 4.02 min, purity >98 %.

### Ligand binding displacement assays

Inhibitor binding was determined using a binding-displacement assay, which measures the ability of inhibitors to displace a fluorescent tracer compound from the ATP binding site of the kinase domain. Inhibitors were dissolved in DMSO and dispensed as 16-point, 2× serial dilutions in duplicate into black multi-well plates (Greiner). Each well contained either 0.5 nM or 1 nM biotinylated kinase domain protein ligated to streptavidin-Tb-cryptate (Cisbio), 12.5 nM or 25 nM Kinase Tracer 236 (ThermoFisher Scientific), 10 mM HEPES pH 7.5, 150 mM NaCl, 2 mM DTT, 0.01 % BSA, 0.01 % Tween-20. Final assay volume for each data point was 5 μL, and final DMSO concentration was 1 %. The kinase domain proteins were expressed in *E. coli* as a fusion with a C-terminal AVI tag (vector pNIC-Bio3, NCBI reference JN792439) which was biotinylated by coexpressed BirA, and purified using the same methods as used previously.^2^ After setting up the assay plate it was incubated at room temperature for 1.5 h and read using a TR-FRET proto Residue ranges were AAK1: 31–396, BMP2K: 38–345, GAK: 12–347, STK16: 13–305 col on a PheraStarFS plate reader (BMG Labtech). The data were normalized to 0 % and 100 % inhibition control values and fitted to a four-parameter dose-response binding curve in GraphPad Software (ver. 7, La Jolla, CA, USA). The determined IC_50_ values were converted into *K*_i_ values using the Cheng–Prusoff equation and the concentration and *K*_D_ values for the tracer (previously determined).

### Competition binding assays

Competition binding assays were performed at DiscoverX as described previously.^30^

### Isothermal Titration Calorimetry

The ITC measurement was performed as described using a NanolTC (TA instruments) at 15 °C in buffer (50 mM HEPES pH 7.5, 500 mM NaCl, 0.5 mM TCEP and 5% Glycerol).^9^ GAK (95 μM) was injected into the cell that contained compound **11** (9 μM). The integrated heat of the titration was calculated and fitted to a single, independent binding model using the software provided by the manufacturer. The thermodynamic parameters ΔH and ΓΔS, as well as the association and dissociation constants (*K*_a_ and *K*_D_) and the stoichiometry were calculated. The data was graphed using GraphPad Prism 7.

### Cellular NanoBRET target engagement assays

Assays were performed essentially as described previously.^22^ In brief: Full-length GAK and RIPK2 ORF (Promega) cloned in frame with a *N*-terminal NanoLuc-fusion and C-terminal NanoLuc-fusion, respectively, were transfected into HEK293T cells and proteins were allowed to express for 20 h. 16-point serially diluted inhibitor and NanoBRET Kinase Tracer K5 (Promega) at 0.5 μM for GAK and NanoBRET Kinase Tracer K4 (Promega) at 0.5 μM for RIPK2, respectively, were pipetted into white 384-well plates (Greiner 781207). The corresponding GAK or RIPK2 transfected cells were added and reseeded at a density of 2 × 10^5^ cells/mL after trypsinization and resuspending in Opti-MEM without phenol red (Life Technologies). The system was allowed to equilibrate for 2 h at 37 °C / 5% CO_2_ prior to BRET measurements. To measure BRET, NanoBRET NanoGlo Substrate + Extracellular NanoLuc Inhibitor (Promega) was added as per the manufacturer’s protocol, and filtered luminescence was measured on a CLARIOstar plate reader (BMG Labtech) equipped with 450 nm BP filter (donor) and 610 nm LP filter (acceptor). Competitive displacement data were then graphed using GraphPad Prism 7 software using the 4-parameter curve fit with the following equation:

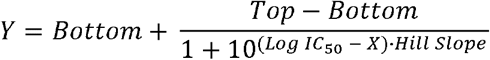

### Prostate cancer cell line viability

Prostate cancer cells (22Rv1, LNCaP, VCaP, PC3, DU145) were obtained from the American Type Culture Collection (Manassas, VA, USA) and cultured in RPMI-1640 medium supplemented with 10% fetal bovine serum and 1% penicillin-streptomycin. LNCaP cells (6000 cells/well) were plated in 96-well plates 24 h prior treatment and treated in triplicate with the designated concentration of compounds. The serial dilutions of compounds were made from 10 mM DMSO stocks in serum-free RPMI-1640. After 48 or 72 h, a 3-(4,5-dimethylthiazol-2-yl)-5-(3-carboxymethoxyphenyl)-2-(4-sulfophenyl)-2H-tetrazolium (MTS) reagent (CellTiter 96^®^ aqueous one solution reagent) (Promega, Madison, USA) was added into each well of the 96-well plate and incubated at 37 °C for 2 h to measure cell viability and inherent cytotoxicity. The absorbance, which is proportional to the number of viable cells, was recorded at 490 nm using VersaMax Microplate Reader (Molecular Devices). Cell viability was normalized to vehicle-treated cells.

### Western blot analysis

Cells were washed with PBS and lysed in RIPA buffer (radioimmune precipitation assay buffer; 50 mM Tris-HCl, 150 mM NaCl, 5 mM EDTA, 1% Triton X-100, 0.1% SDS, and 1% sodium deoxycholate) and a mixture of protease inhibitors (Sigma). The isolated total protein (20 μg) was run on 4–10 % pre-cast Bis-Tris gels (Invitrogen) and electrotransferred to a 0.2-μm polyvinylidene difluoride (PVDF) membrane. Membranes were blocked for 30 min in StartingBlock (PBS) blocking buffer (Pierce) and probed with antibodies for PARP (9532S, Cell Signaling Technology), phosphor-histone H3 (3377S, Cell Signaling Technology), AR (N-20, sc-816, Santa Cruz Biotechnology Inc.), and β-actin (4970S, Cell Signaling Technology). The PARP antibody can detect both full-length (116 kDa) and cleaved PARP (89 kDa). β-actin was used as a loading control. Detection was performed using ECL HRP-linked anti-mouse or anti-rabbit IgG (GE Healthcare) and ECL™ Prime (GE Healthcare). Blots were imaged on a ChemiDoc MP Imaging System (Bio-Rad).

### Mass Spectrometry and LC method

Samples were analyzed with a hybrid LTQ FT (ICR 7T) (ThermoFisher, Bremen, Germany) mass spectrometer coupled with a Waters Acquity H-class liquid chromatograph system. Samples were introduced via an electrospray source at a flow rate of 0.6 mL/min. Electrospray source conditions were set as: spray voltage 4.7 kV, sheath gas (nitrogen) 45 arb, auxiliary gas (nitrogen) 30 arb, sweep gas (nitrogen) 0 arb, capillary temperature 350 °C, capillary voltage 40 V and tube lens voltage 100 V. The mass range was set to 150-2000 *m*/*z*. All measurements were recorded at a resolution setting of 100,000.

Separations were conducted on a Waters Acquity UPLC BEH C18 column (2.1 x 50 mm, 1.7 μm particle size). LC conditions were set at 100 % water with 0.1 % formic acid (A) ramped linearly over 9.8 min to 95 % acetonitrile with 0.1 % formic acid (B) and held until 10.2 min. At 10.21 min the gradient was switched back to 100 % A and allowed to re-equilibrate until 11.25 min.

Xcalibur (ThermoFisher, Breman, Germany) was used to analyze the data. Solutions were analyzed at 0.1 mg/mL or less based on responsiveness to the ESI mechanism. Molecular formula assignments were determined with Molecular Formula Calculator (v 1.2.3). Low-resolution mass spectrometry (linear ion trap) provided independent verification of molecular weight distributions. All observed species were singly charged, as verified by unit *m*/*z* separation between mass spectral peaks corresponding to the ^12^C and ^13^C^12^C_c-1_ isotopes for each elemental composition.

